# TREM2 drives monocyte-derived macrophage responses to *Cryptococcus neoformans*

**DOI:** 10.64898/2026.06.10.731331

**Authors:** Adiza Abass, Alison Ricafrente, Apurwa Trivedi, Amy Vichaidit, Aelita Arshakyan, Sreemoyee Acharya, Lena J. Heung

## Abstract

*Cryptococcus neoformans* is an opportunistic fungus that causes pulmonary and central nervous system infections after entry into the lungs. We previously established that signaling through the adapter protein DAP12 inhibits the antifungal response of monocyte-derived macrophages and worsens the survival of mice after *C. neoformans* infection. However, the molecular mechanisms by which DAP12 signaling is initiated during cryptococcosis remain inadequately characterized. In this study, we identify triggering receptor expressed on myeloid cells 2 (TREM2) as a DAP12-associated receptor that is induced on murine monocytes and interstitial macrophages in the lungs in response to *C. neoformans* infection. TREM2 subsequently represses fungal uptake and M1 polarization by monocyte-derived macrophages. Using an *in vitro* binding assay, we find that both murine and human TREM2 can directly bind to *C. neoformans* and that the absence of the cryptococcal cell wall antigen β-1,6-glucan disrupts these interactions. Overall, our findings suggest that the TREM2-DAP12 pathway plays an important inhibitory role in the host immune response to cryptococcal infection by impeding macrophage activation and phagocytosis of fungal cells. We also establish TREM2 as a receptor involved in direct fungal sensing of *C. neoformans*.

## Introduction

*Cryptococcus neoformans* is an environmental fungus that can infect the lungs and central nervous system and is the leading cause of death in immunocompromised patients, particularly those with AIDS, cancer, or organ transplantation (1). Yet, our understanding of how *C. neoformans* is sensed by and interacts with the host immune system upon initial entry into the lungs is still not well defined (2).

In previous work, we found that activation of the mammalian DNAX*-*activating protein of 12 kDa (DAP12) suppresses the monocyte-derived macrophage response to *C. neoformans* and worsens outcomes in a murine model of infection (3). DAP12 is a transmembrane signaling adapter that pairs to several cell surface receptors on myeloid cells and natural killer (NK) cells (4–8). In this study, we evaluated the potential role of triggering receptor expressed on myeloid cells 2 (TREM2), a known DAP12-associated receptor, in the initiation of DAP12 signaling by monocyte-derived macrophages during *C. neoformans* infection. TREM2 is a transmembrane cell surface receptor with an immunoglobulin-like extracellular domain that is found on myeloid cells (9–11) and is known to bind to a variety of molecules and antigens on microbial and mammalian cells (12–14). TREM2 on microglia plays an important role in several neurodegenerative diseases (12). Additionally, TREM2 has been studied in macrophages during bacterial, viral, and parasitic infections and can induce diverse cellular effects (15–22). Of note, TREM2 promotes the proliferation and survival of M2 macrophages during bacterial and viral infections (20, 23), limits the anti-mycobacterial activity of macrophages (24) and inhibits the production of inflammatory cytokines (9, 13).

In this study, we demonstrate that after *C. neoformans* infection, TREM2 is upregulated on lung monocytes and macrophages and inhibits M1 polarization, TNF secretion, and phagocytosis of *C. neoformans* by monocyte-derived macrophages. We also discover that TREM2 is a pattern recognition receptor for *C. neoformans*. Thus, by inducing TREM2-DAP12 signaling in macrophages, *C. neoformans* can suppress key elements of the antifungal response by the host.

## Results

### Inflammatory monocytes in the lungs of *C. neoformans*-infected mice express DAP12-associated receptors, including TREM2

To identify potential receptors that initiate DAP12 signaling during *C. neoformans* infection, we utilized our previously generated RNA-seq data on CCR2^+^Ly6C^hi^ inflammatory monocytes sorted from the lungs of CCR2-GFP reporter mice (25) on Day 10 post-infection (p.i.) with *C. neoformans* strain H99 (GEO Series accession number GSE122765) (26). We analyzed the expression of known DAP12-associated receptors by these inflammatory monocytes and found that among the transcripts identified was triggering receptor expressed on myeloid cells 2 (TREM2) (Fig. 1A). TREM2 is a member of the immunoglobulin superfamily that has been shown to limit macrophage responses to bacterial and viral infections (20, 24). Although TREM2 was not the most highly expressed DAP12-associated receptor, its ability to inhibit macrophage activity is similar to the phenotype that we observed with DAP12 in our previous work (3).

**Fig. 1:**
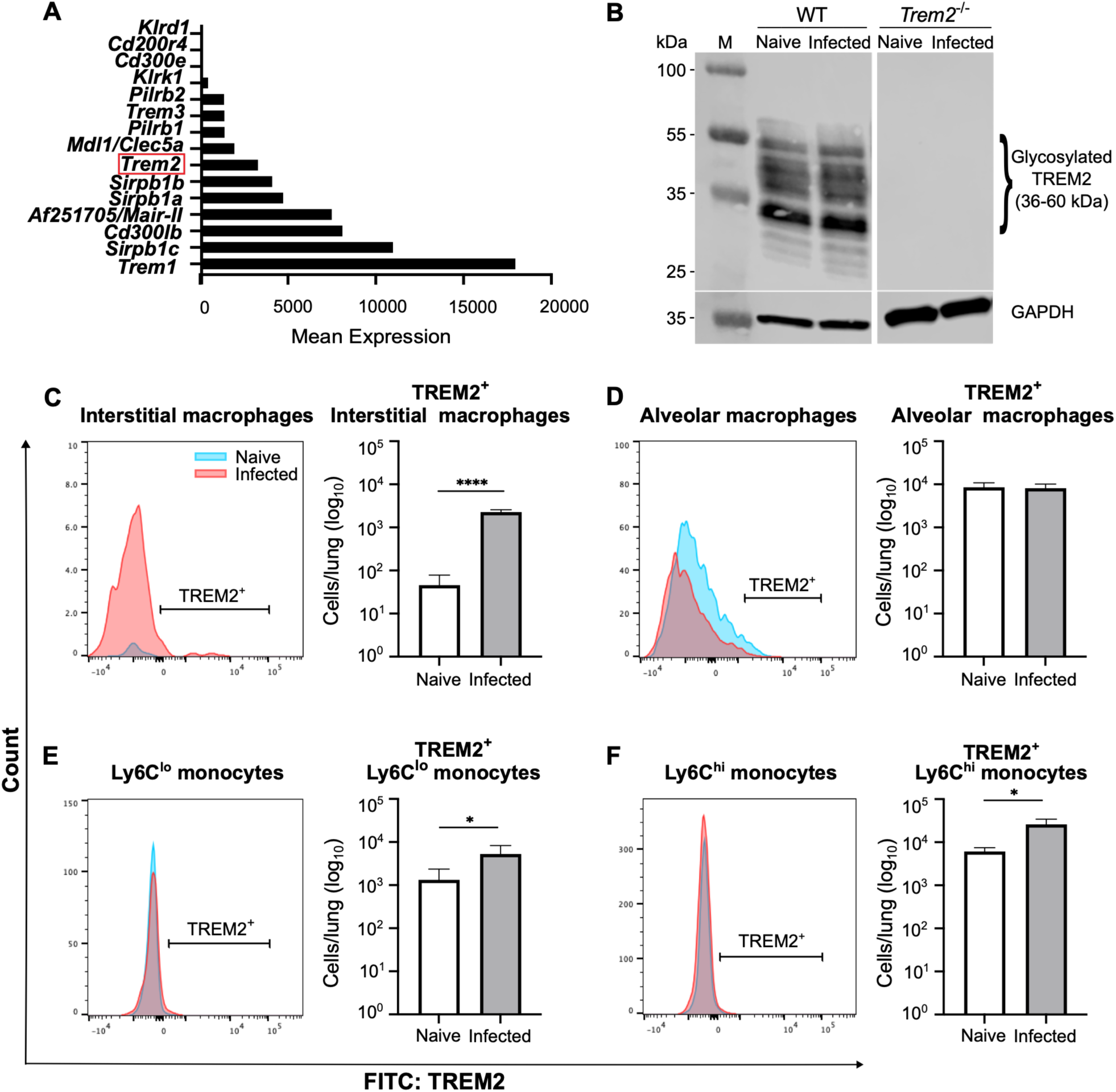
*C. neoformans* induces the expression of TREM2 on monocytes and macrophages. (A) Mean expression of DAP12-associated receptors by inflammatory CCR2^+^Ly6C^hi^ monocytes at Day 10 p.i. with *C. neoformans* from previously generated RNA-seq data (NCBI GEO Series accession GSE122765) (26). (B) Immunoblot of TREM2 expression in the lysate of WT and *Trem2*^-/-^ BMDM, either naive or challenged with *C. neoformans* for 40 min at MOI 1:40, with GAPDH as a loading control. (C-F) TREM2 expression on monocytes and macrophages from the lungs of naive and *C. neoformans-*infected WT mice on Day 7 p.i. were analyzed via flow cytometry. Representative histograms are shown on the left of each panel. Blue = naive, Pink = infected. Quantification of the number of TREM2^+^ cells in each subpopulation is shown in the bar graphs on the right of each panel. White = naive, Gray = infected. Data are from three independent experiments (*n* = 5 total mice per group). *, *P* < 0.05; ****, *P* < 0.0001 by *t*-test.

Together, these findings suggested that TREM2 may pair with DAP12 on monocytes and monocyte-derived macrophages (moMacs) during *C. neoformans* infection.

### *C. neoformans* induces TREM2 expression on monocyte-derived macrophages

To evaluate if TREM2 protein is expressed by moMacs during infection, we first utilized C57BL/6J wild type (WT) and *Trem2*^-/-^ bone marrow-derived macrophages (BMDM).

Immunoblot analysis of whole cell lysates of naive BMDM or BMDM challenged with *C. neoformans* strain H99 revealed that BMDM express TREM2 under both steady state and infected conditions (Fig. 1B); the molecular weight of glycosylated murine TREM2 ranges from 36-60 kDa (27). As expected, there was no TREM2 detected in naive or infected *Trem2^-/-^* BMDM (Fig. 1B). Thus, TREM2 is expressed by WT moMacs *in vitro*.

We then investigated the expression of TREM2 by lung monocytes and macrophages in our mouse model of infection and found that *C. neoformans* significantly increases the expression of TREM2 on interstitial macrophages in WT mice by Day 7 p.i. compared to naive mice (Fig. 1C). We also observed increases in TREM2 expression on Ly6C^lo^ and Ly6C^hi^ monocytes but not on alveolar macrophages from infected mice on Day 7 p.i. (Fig. 1D through F). TREM2 induction on interstitial macrophages persists through Day 21 p.i., with no changes on monocytes or alveolar macrophages at the same timepoint (Fig. S1). Since interstitial macrophages are predominantly monocyte-derived cells (28), these results further indicate that TREM2 has a specific role on this cellular subset during cryptococcosis.

### TREM2 suppresses monocyte-derived macrophage antifungal responses

To determine the role of TREM2 in the moMac response to *C. neoformans*, we challenged WT and *Trem2^-/-^* BMDM with a fluorescent *C. neoformans* strain H99-GFP (29). We used flow cytometry to identify BMDM that take up the GFP^+^ fungus and found that *Trem2^-/-^*BMDM have an approximately two-fold increase in uptake of the fungus compared to WT BMDM at two different multiplicities of infection (MOI) (Fig. 2A). There was no significant difference in fungal killing as measured by CFU (Fig. 2B). However, *Trem2^-/-^* BMDM have increased secretion of the pro-inflammatory cytokine TNF compared to WT BMDM (Fig. 2C). This increased fungal uptake and TNF secretion by

**Fig. 2:**
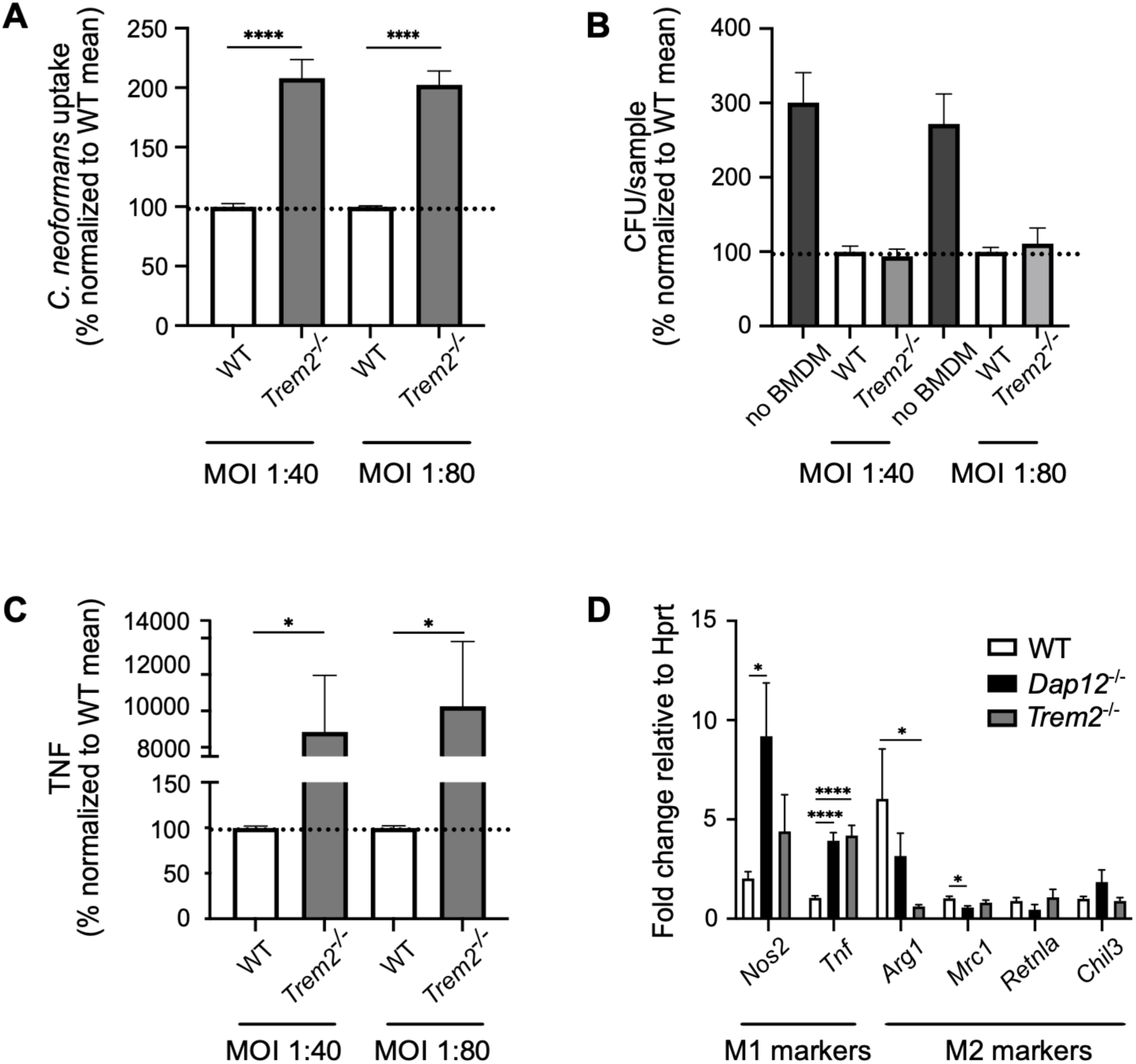
TREM2 regulates monocyte-derived macrophage responses to *C. neoformans*. BMDM from WT mice (white bars), *Trem2*^-/-^ mice (light gray bars) and *Dap12*^-/-^ mice (black bars) were incubated with GFP-expressing *C. neoformans* for 24 hours. (A) Fungal uptake was measured as the percentage of BMDM associated with the fungal GFP signal by flow cytometry (WT mean at MOI 1:40 = 2.4% of total macrophages containing GFP^+^ fungal cells and WT mean at MOI 1:80 = 0.7% of total macrophages containing GFP^+^ fungal cells). (B) Fungal killing was measured by plating CFU (WT mean at MOI 1:40 = 232,750 CFU and WT mean at MOI 1:80 = 134,583 CFU). For reference, growth of *C. neoformans* in the absence of BMDM is shown (No BMDM, dark gray bars). (C) TNF secretion was measured by ELISA of culture supernatants (WT mean at MOI 1:40 = 2.5 pg/mL and WT mean at MOI 1:80 = 2.4 pg/mL). (D) Macrophage polarization markers were assessed by quantitative RT-PCR. Data are from two to three independent experiments (*n* = 3 replicates per group). *, *P* < 0.05; ***, *P* < 0.001 by *t-*test (panels A-C) and one-way ANOVA (panel D).

*Trem2*^-/-^ BMDM recapitulates the responses of *Dap12^-/-^*BMDM to *C. neoformans* in our previous work (3). Furthermore, quantitative RT-PCR analysis showed significant upregulation in the transcription of *Tnf*, an M1 macrophage marker, in both *Trem2*^-/-^ and *Dap12*^-/-^ BMDM and downregulation of the M2 macrophage markers *Arg1* and *Mrc1* in *Trem2^-/-^*and *Dap12*^-/-^ BMDM, respectively, when compared to WT BMDM (Fig. 2D).

There was also increased expression of the M1 marker *Nos2* in *Dap12^-/-^* BMDM relative to WT BMDM (Fig. 2D). Thus, both TREM2 and DAP12 skew BMDM polarization towards an M2 phenotype. Overall, these results indicate that TREM2-DAP12 signaling inhibits fungal uptake and pro-inflammatory responses by moMacs during *C. neoformans* challenge.

Next, we investigated the role of TREM2 *in vivo* by administering *C. neoformans* H99 intratracheally (i.t.) to WT and *Trem2^-/-^*mice. No significant differences in lung fungal burden on Days 7 and 14 p.i. or in overall survival between the two mouse strains were observed (Fig. 3A and B). Cytokine profiling of whole lung homogenates showed no significant differences between *Trem2^-/-^* and WT mice (Fig. S2). Total numbers of innate immune cells remained comparable in WT and *Trem2^-/-^* mice, except for a small but transient decrease in CD103^+^ conventional DCs (cDCs) (Fig. S3).

**Fig. 3:**
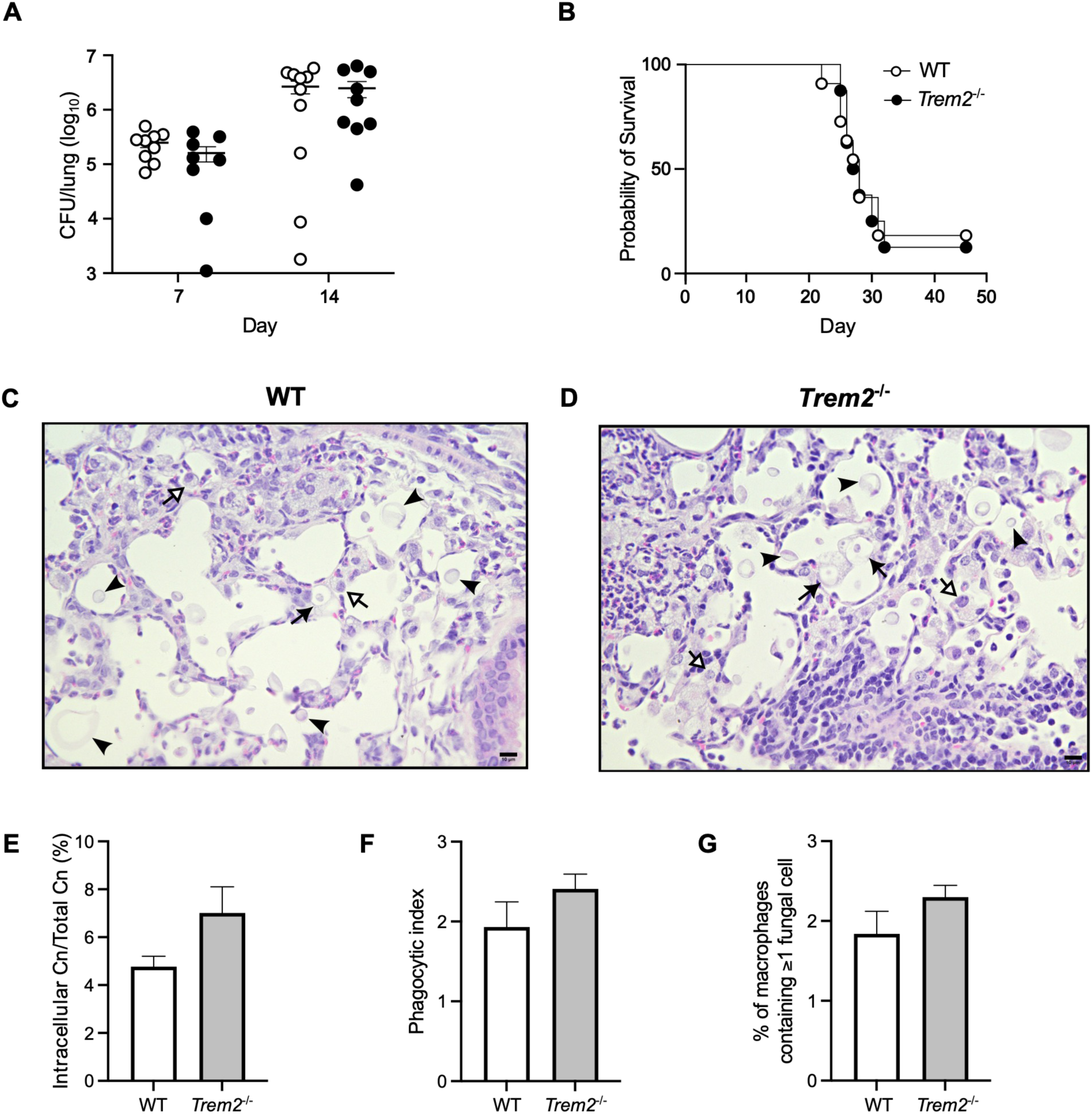
TREM2 regulates macrophage uptake of *C. neoformans* in the lungs. (A) Lung fungal burden in WT mice (white circles) and *Trem2^-^*^/-^ mice (black circles) on Days 7 and 14 p.i. Data are from two independent experiments (*n* = 8-10 total mice per group). (B) Kaplan-Meier survival curve of infected WT and *Trem2*^-/-^ mice. Data are pooled from three independent experiments (*n* = 19-21 total mice per group). (C-D) Representative images of lung tissue sections from WT (C) and *Trem2*^-/-^ (D) mice on Day 7 p.i. stained with H&E. Scale bar = 10 µM at 40X magnification. Arrowhead = extracellular fungal cell, black arrow = macrophage containing fungal cell, white arrow = macrophage. (E-G) Quantification of histology findings including the percentage of *C. neoformans* cells that are intracellular (E), phagocytic index (the number of engulfed fungal cells per 100 macrophages) (F), and the percentage of macrophages containing ≥1 *C. neoformans* cell (G). For results in E through G, two entire lung slices were evaluated from each of 3 total mice per group. Data were analyzed using *t*-test (A, E-G) and Mantel-Cox (B).

To determine whether TREM2 regulates phagocytosis of *C. neoformans in vivo*, as we observed *in vitro* (Fig. 2A), analysis of H&E-stained lung sections from WT and *Trem2^-/-^*mice on Day 7 p.i. was performed (Fig. 3C through F; Fig. S4). There was no significant difference in overall lung pathology (Fig. S4), but closer inspection suggested that *C. neoformans* was found within *Trem2*^-/-^ lung macrophages more frequently than WT macrophages (Fig. 3C and D). This observation was supported by trends in the quantitation of our histology findings, including the percentage of *C. neoformans* cells that are found within macrophages (Fig. 3E), the phagocytic index (Fig. 3F), which is the total number of engulfed fungal cells per 100 macrophages, and the percentage of macrophages that have ingested at least one fungal cell (Fig. 3G). Thus, in line with our *in vitro* BMDM data, TREM2 may limit the uptake of *C. neoformans* by pulmonary macrophages *in vivo*. Although the loss of TREM2 does not appear to be sufficient to reverse overall infectious outcomes, TREM2 may still play a key role in initial host sensing of *C. neoformans* and subsequent inhibition of phagocytosis and M1 polarization by moMacs.

### TREM2 binds directly to *C. neoformans*

Since TREM2 regulates *C. neoformans* uptake by moMacs, we wondered if TREM2 directly interacts with the fungus. TREM2 is known to bind to a variety of endogenous and microbial ligands (12–14), so we performed binding assays using soluble, recombinant TREM2-Fc proteins consisting of the extracellular domain of either mouse or human TREM2 conjugated to human Fc (14). We found that both mouse and human TREM2-Fc bind to encapsulated *C. neoformans* H99, and that they bind even more avidly to an acapsular mutant of H99 called cap59 (30) (Fig. 4). These results suggest that TREM2 interacts with a fungal cell wall antigen that is more exposed on the acapsular mutant. TREM2-Fc did not bind to the unencapsulated yeast *Candida albicans* in our assay (Fig. 4), as was previously shown (14). Additionally, TREM2-Fc did not bind to *Cryptococcus gattii* (Fig. 4), an encapsulated species that causes infections similar to *C. neoformans*, but has distinct differences in epidemiology (31). Together, these findings indicate that TREM2 binds directly to a cell wall antigen of *C. neoformans* and that this interaction is unique to *C. neoformans* compared to several other pathogenic yeasts.

**Fig. 4:**
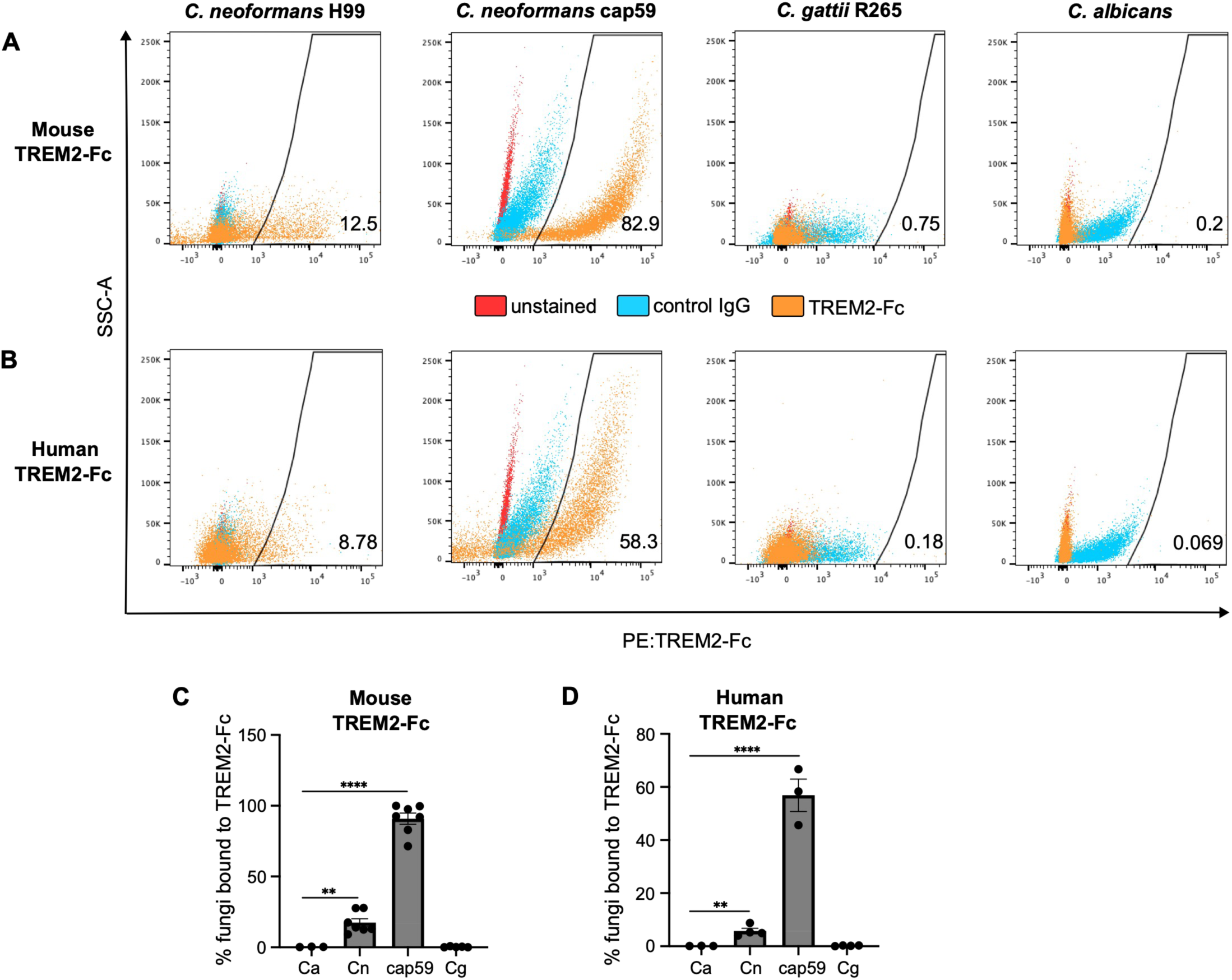
TREM2 binds uniquely to *C. neoformans*. (A-D) Fungal cells were incubated with either a recombinant TREM2-Fc protein, that fuses the murine (A, C) or human (B, D) TREM2 extracellular domain with the Fc domain of human IgG, or with purified human IgG as a control. Binding was measured using a PE-conjugated secondary antibody against the Fc domain of human IgG and flow cytometry to identify fungal cells associated with the PE signal. Shown are representative dot plots of unstained (red), human IgG-bound (blue), and TREM2-Fc bound (orange) cells of *C. neoformans* H99 (Cn), an acapsular *C. neoformans* strain cap59, another encapsulated species *C. gattii* (Cg), and the yeast *Candida albicans* (Ca) as a negative control (A, B). Quantification of all results are shown in the bar graphs (C, D). Statistics were measured in reference to Ca. Data were pooled from three to six independent assays *(n* = 2 replicates per fungal strain per experiment). **, *P* < 0.01; ****, *P* < 0.0001 by one-way ANOVA.

### TREM2 binding to *C. neoformans* is dependent on β-1,6-glucan

To identify the cell wall component of *C. neoforman*s that interacts with TREM2, we tested the binding of mouse and human TREM2-Fc to *C. neoformans* mutants in which the enzymes involved in the synthesis of key cryptococcal cell wall antigens α-1,3-glucan, chitosan, and β-1,6-glucan have been deleted (Fig. 5A; Table S1). There was no significant difference in the binding of TREM2-Fc to the *ags1*Δ strain, which lacks α-1,3- glucan, and its parental strain H99R (referred to as WT) (Fig. 5B and C) (32, 33). We next examined *C. neoformans* strains with defects in chitosan synthesis. *C. neoformans* generates chitin through the action of chitin synthase 3 (CHS3); this chitin is then converted to chitosan via chitosan deacetylase (CDA) enzymes (34, 35). The *cda1*Δ*cda2*Δ and *cda1*Δ*cda2*Δ*cda3*Δ strains (35), which have moderate and severe loss of chitosan, respectively, exhibit a significant loss of mouse TREM2-Fc binding compared to the parental strain KN99 (referred to as WT) (Fig. 5D) (36) but no changes with human TREM2-Fc binding (Fig. 5E). Interestingly, the *chs3*Δ strain, that has a defect in the synthesis of chitin which results in a significant loss of chitosan (34, 35), showed no changes in mouse or human TREM2-Fc binding versus WT (Fig. 5D and E). Finally, we tested the *kre5*Δ and *skn1*Δ*kre6*Δ strains (37) that have defects in the production of β-1,6-glucan, the predominant β-glucan in the *C. neoformans* cell wall (38), and found that both mutant strains have decreased binding of mouse and human TREM2-Fc (Fig. 5G and H). Thus, our studies identify TREM2 as a new fungal pattern recognition receptor that may bind to β-1,6-glucan in the cell wall of *C. neoformans* to impede macrophage activation and uptake of fungal cells during infection.

**Fig. 5:**
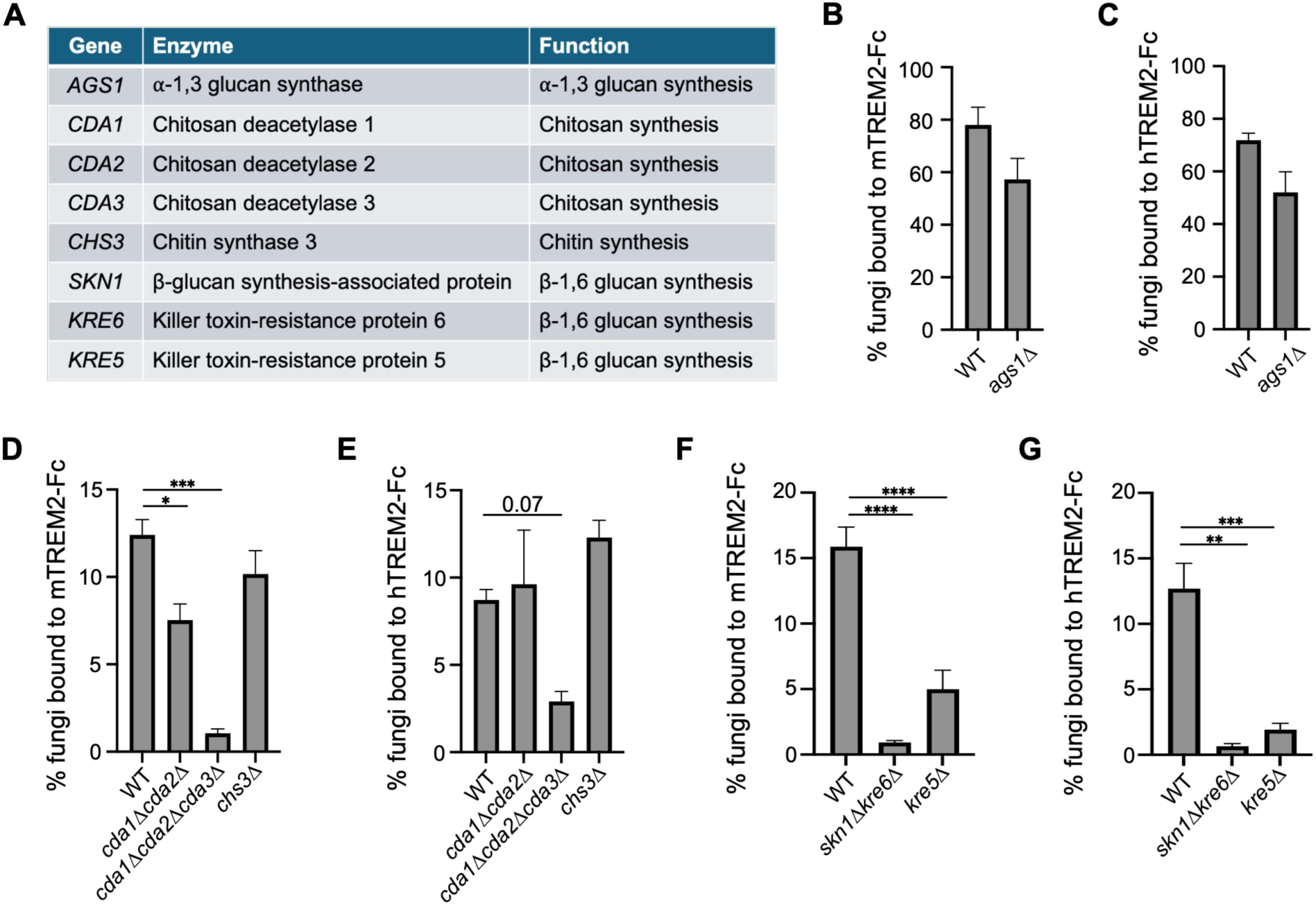
TREM2 binds to β-1,6-glucan on the cell wall of *C. neoformans*. (A) Key *C. neoformans* genes and their corresponding enzymes that are involved in cell wall antigen synthesis. (B-G) Quantification of fungal cells bound to either mouse (m) or human (h) recombinant TREM2-Fc protein. The bar graphs compare WT *C. neoformans* H99R versus the ⍺-1,3-glucan-deficient strain *ags1*Δ (B, C); WT *C. neoformans* KN99 versus chitosan-deficient strains *cda1*Δ*cda2*Δ, *cda1*Δ*cda2*Δ*cda3*Δ, and *chs3*Δ (D, E); and WT *C. neoformans* H99 versus β-1,6-glucan-deficient strains *skn1*Δ*kre6*Δ and *kre5Δ* (F, G). Data were pooled from two to five independent experiments (*n* = 2-3 replicates per fungal strain). Significance was measured in reference to the WT strain. *, *P* < 0.05; **, *P* < 0.01; ***, *P* < 0.001; ****, *P* < 0.0001 by *t*-test (B, C) and one-way ANOVA (D-G).

## Discussion

In this study, we found that TREM2, a DAP12-associated receptor, plays a novel role in the recognition of *C. neoformans* and the subsequent macrophage response to the fungus (Fig. 6). The TREM2-DAP12 signaling pathway can be either protective or detrimental to the host depending on the disease model (6, 39). In the case of *C. neoformans* infection, we previously demonstrated that DAP12 impairs M1 polarization as well as fungal uptake and killing by moMacs, resulting in worse infectious outcomes (3). Here, we show that DAP12 signaling initiated by TREM2 specifically regulates macrophage polarization and phagocytosis in response to *C. neoformans* infection. Similarly, TREM2 impedes M1 macrophage polarization during mycobacterial infection (24, 40) and inhibits macrophage uptake of the bacterium *Streptococcus pneumoniae* (18). A clinical study also identified a positive correlation between TREM2 and the expression of M2-associated macrophage markers in the lungs of patients with pulmonary tuberculosis compared with healthy controls (41). However, a mechanistic role for TREM2 during fungal infection has not been previously described. Daws et al. showed that TREM2 can bind to the yeast *Candida guilliermondii* (14), but the functional consequences for the host response to this yeast remain unclear.

**Fig. 6:**
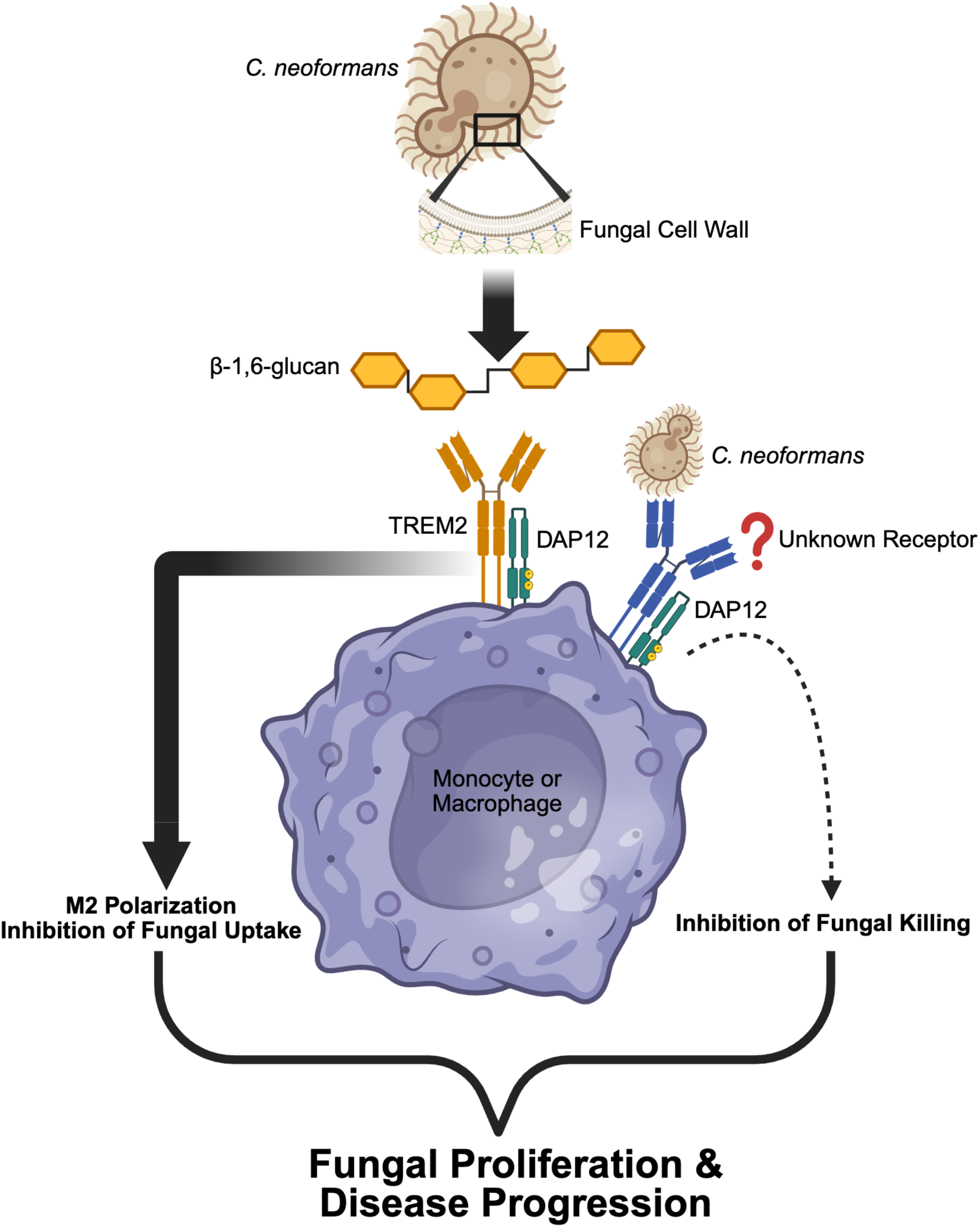
Proposed model of TREM2-DAP12 signaling during infection with *C. neoformans.* β-1,6-glucan within the cell wall of *C. neoformans* is recognized by TREM2 on monocytes or macrophages and induces TREM2-DAP12 signaling. This signaling drives M2 polarization and reduces fungal uptake by monocyte-derived macrophages. Concurrently or sequentially, we hypothesize that DAP12 engages another yet unknown receptor to also inhibit fungal killing, which is regulated by DAP12 based on our previous work (3). In concert, these DAP12-mediated pathways can facilitate fungal proliferation and disease progression during *C. neoformans* infection. (Created in BioRender. Ricafrente, A. (2026) https://BioRender.com/vnn9cbd)

It has been proposed that DAP12 and other immunoreceptor tyrosine-based activation motif (ITAM)-containing signaling adapters can induce a wide range of cellular responses, including both positive and negative regulatory effects, depending on ligand affinity or avidity to the associated receptor (6, 42). TREM2 is a promiscuous receptor that binds to an equally wide variety of endogenous and exogenous ligands (12, 43).

Commonly, TREM2 ligands are lipids, but protein ligands have been described, including the lectin galectin-3 (44). Our findings expand this repertoire to include β-1,6-glucan, a polysaccharide component of the fungal cell wall (45). Interestingly, β-1,6-glucan is the predominant glucan in the cell wall of *C. neoformans*, not β-1,3-glucan like many other fungi (38, 46). Particles containing zymosan, a cell wall preparation from the yeast *Saccharomyces cerevisiae* that is predominantly composed of β-1,3-glucan, do not bind TREM2 (47). It is not clear why we do not observe binding of TREM2 to *C. gattii*, but differences in β-1,6-glucan content or exposure between the *Cryptococcus* species have not yet been evaluated. In our study, we also determined that human TREM2 binds to *C. neoformans*; TREM2 and its family of receptors are known to be evolutionarily conserved (43). Structural studies indicate that there are at least three different regions of the immunoglobulin domain of human TREM2 that may serve as binding sites for multiple ligands (48–51). Additional work will be needed to determine the structural nature of the interaction between TREM2 and β-1,6-glucan.

While TREM2 serves as a phagocytic receptor for many bacteria (40, 47, 52), our data indicate that TREM2 is not a phagocytic receptor for *C. neoformans*. Rather, engagement of TREM2 by the fungus likely inhibits other pathways or mechanisms involved in phagocytosis. TREM2-DAP12 has previously been shown to inhibit signaling through Toll-like receptors (TLRs) (9, 53), innate immune receptors that can regulate different facets of the phagocytic process by macrophages (54). TLR agonists can enhance phagocytosis of *C. neoformans* by microglia, the tissue-resident macrophages of the central nervous system (55). However, the role of TLRs in macrophage uptake of *C. neoformans* in other compartments remains unclear. Of note, TLR4 has been reported to inhibit macrophage phagocytosis of *C. neoformans in vitro* (56), though the contributions of TLR4, as well as TLR2, to overall outcomes in cryptococcosis remain controversial (57–59). TLR9, an intracellular receptor, regulates macrophage polarization and is essential for host defense (60–63), but it has not been implicated in phagocytosis during *C. neoformans* infection. Thus, the likelihood that TREM2-DAP12 inhibits macrophage uptake of *C. neoformans* via TLR regulation is low, although this pathway could modulate macrophage polarization. In a respiratory infection model with *S. pneumoniae*, TREM2 suppresses C1q complement protein, thereby reducing opsonization and subsequent phagocytosis by macrophages (18). Whether TREM2 regulates phagocytosis of *C. neoformans* through these or other indirect mechanisms will need to be addressed in future studies.

As a facultative intracellular pathogen, it may seem counterintuitive that *C. neoformans* would inhibit its own uptake via TREM2. This mechanism may serve as a means of bet-hedging to enhance overall survival. Fungal cells that remain extracellular due to TREM2-mediated inhibition of phagocytosis are less likely to be killed by M1-polarized macrophages; concurrently, fungal cells phagocytosed through TREM2-independent pathways may encounter TREM2-induced M2-polarized macrophages, which are typically permissive for intracellular survival and proliferation (64). The persistent induction of TREM2 on interstitial macrophages that we observed could sustain this strategy over the course of infection.

In summary, TREM2 is upregulated on moMacs during *C. neoformans* challenge, binds directly to *C. neoformans* through the fungal cell wall antigen β-1,6-glucan, and inhibits M1 polarization of these macrophages and their phagocytosis of fungal cells. To our knowledge, this study is the first to demonstrate a functional role for TREM2 in host interactions with a fungal pathogen, highlighting the receptor’s capacity to recognize diverse ligands and regulate multiple cellular functions. Notably, the phenotypes observed in the *Trem2*^-/-^ mice do not recapitulate all the effects seen in the absence of DAP12 (3), suggesting that additional receptors act concurrently or sequentially with TREM2 to signal through DAP12. Thus, DAP12-coupled receptors remain an important area of investigation for understanding host interactions with *C. neoformans*.

## Materials and Methods

### Chemicals and reagents

All chemicals and reagents were obtained from Thermo Fisher Scientific unless otherwise specified.

### Fungal and mouse strains

*Cryptococcus neoformans* strain H99 #4413 (referred to as H99, provided by Joseph Heitman at Duke University), *C. neoformans* strain H99-GFP (provided by Robin May at University of Birmingham) (29), *Cryptococcus gattii* strain R265 (ATCC MYA-4093), and *Candida albicans* strain SC5314 (provided by Tobias Hohl at Memorial Sloan Kettering Cancer Center) were grown from frozen glycerol stocks on Sabouraud dextrose agar plates (SAB) at 37°C for 72 hours. *C. neoformans* strain cap59 (provided by Joseph Heitman at Duke University) (30) was grown from frozen glycerol stocks on YPD (1% yeast extract, 2% peptone, 2% dextrose) agar plates at 37°C for 72 hours. Single colonies were then inoculated into YPD broth and grown overnight on a 37°C shaker at 250 rpm, unless otherwise noted. *C*. *neoformans* cell wall mutant strains and the parental strains KN99 (36) and H99R [serotype A *MATα ura5*] (33) (see Table S1) were generously provided by Tamara Doering (Washington University in Saint Louis) and Jennifer Lodge (Duke University). These mutant strains were grown from frozen glycerol stocks on SAB plates at 30°C for 72 hours, followed by growth in YPD broth on a 30°C shaker at 300 rpm for 48 hours. For TREM2 binding assays, all strains were cultured in the same manner as the mutant strains. *Trem2^-/-^* (strain #027197) and C57BL/6J (strain #000664) mice were obtained from Jackson Laboratories. *Dap12^-/-^* mice were obtained from Taconic (strain #12665)(65).

### Ethics statement

All animal studies were performed with approval from the Cedars-Sinai Institutional Animal Care and Use Committee (IACUC) under protocol 8429 and were compliant with all applicable provisions established by the Animal Welfare Act and the Public Health Services Policy on Humane Care and Use of Laboratory Animals.

### RNA-seq analysis

Expression of DAP12-associated receptors by CCR2^+^Ly6C^hi^ monocytes was evaluated using RNA-seq data previously generated (26) and publicly accessible through NCBI Gene Expression Omnibus (GEO) Series accession number GSE122765 (https://www.ncbi.nlm.nih.gov/geo/query/acc.cgi?acc=GSE122765).

### Bone marrow-derived macrophage studies

Bone marrow-derived macrophages were generated using supernatant from L929 cells, as previously described (3, 66). Overnight cultures of *C. neoformans* H99-GFP were opsonized with 10 µg/mL mouse anti-glucuronoxylomannan monoclonal antibody 18B7 antibody (Millipore Sigma) in DMEM media for 1 hour at room temperature (RT) on a shaker. BMDM were challenged with the opsonized fungal cells at multiplicities of infection (MOI) of 1:40 or 1:80 for 24 h at 37°C, 5% CO2 in DMEM supplemented with 10% FBS media. Mouse TNF cytokine secretion was measured by ELISA (Invitrogen) on cell culture supernatant using a Varioskan Lux Multimode Microplate Reader. Fungal killing was analyzed by plating CFU from infected BMDM lysed in sterile water. Uptake of fungal cells by BMDM was measured by flow cytometry on an LSRFortessa (BD), and data were analyzed using FlowJo software (BD Biosciences). For quantitative RT-PCR, total RNA was extracted from cells using TRIzol LS reagent, cDNA was synthesized using a High-Capacity RNA to cDNA Kit (Applied Biosystems), and analysis was performed on a ViiA 7 Real-Time PCR System (Applied Biosystems) using TaqMan Fast Advanced Master Mix and TaqMan Gene Expression Assays including Arg1 (Mm00475988_m1), Mrc1 (Mm01329362_m1), Retnla/Fizz1 (Mm00445109_m1), Hprt (Mm03024075_m1), Chil3/Ym1 (Mm00657889_mH), Nos2 (Mm00440502_m1), and Tnf (Mm00443258_m1).

### Immunoblot assay

BMDM were generated from C57BL/6J (WT) and *Trem2*^-/-^ mice and serum starved for 14 hours prior to challenge with opsonized *C. neoformans* H99 for 40 min at MOI 1:40. Proteins were extracted from infected and uninfected (naive) WT and *Trem2^-/-^* BMDM using NP-40 lysis buffer (Millipore Sigma) supplemented with 1X Halt^TM^ Protease and Phosphatase Inhibitor Cocktail on ice. Cell debris was cleared by centrifugation at 150,000 rpm for 5 min. The protein concentration of the cell lysates was analyzed using a Pierce^TM^ BCA Protein assay kit following the manufacturer’s protocol. The cell lysates were then mixed with SDS loading buffer and boiled at 100°C for 10 min. Samples of 30 µg total protein were separated on a Novex^TM^ 10-20% Tris-Glycine Plus Wedge Well^TM^ gel using a mini gel tank and transferred onto a polyvinylidene fluoride membrane (Invitrogen) with an iBlot2 Gel Transfer Device (Life Technologies). The membranes were blocked in 3% BSA/0.05% Tween-20/Tris-buffered saline (TBS) at 4°C for 1 hour and probed with primary antibodies (anti-TREM2 1:1000, Cell Signaling Technologies, 59621; GAPDH 1:2000, Cell Technologies, 2118S) in TBS overnight at 4°C on a shaker. The membranes were then washed and incubated with horseradish peroxidase-conjugated anti-rabbit IgG secondary antibody (1:2000, Cell Signaling Technologies, 7074) at RT for 1 hour. Immunoreactive proteins were detected by chemiluminescence (Supersignal^TM^ West Pico Plus Chemiluminescent Substrate) on a Molecular Imager Gel Doc XR+ (Bio-Rad).

### Infection and analysis of mice

Mice were infected with 10^3^ *C. neoformans* intratracheally as previously described (26, 67). Lungs were collected and homogenized using a gentleMACS™ Octo Dissociator (Miltenyi Biotec). Specifically, to generate single cell suspension, lung samples were homogenized in 1X PBS containing 2.31 mg/mL Collagenase Type 4 (Worthington Biochemical Corporation), 100 mg/mL DNAse I grade II (Roche) and 0.5% Bovine Serum Albumin (BSA) in gentle MACS C tubes using the “Mouse Spleen 1” program. This was followed by a 45 min incubation at 37°C on a shaker prior to another round of homogenization with the “Mouse Lung 2” program. Serial dilutions of the homogenates were plated on SAB plates to measure fungal burden by counting CFU. For survival studies, infected mice were monitored for signs of illness and distress and then euthanized in accordance with protocols approved by the Cedars-Sinai IACUC.

### Flow cytometry

Whole lung single cell suspensions were prepared as described in “Infection and analysis of mice” above. Red blood cells (RBC) in the lung samples were lysed with 1X RBC lysis buffer (TNB-4300-L100, Cytek) for 5 min. Then, the samples were washed with 0.5% BSA in PBS and passed through a 100 µM cell strainer. Total cells in each sample were counted with a hemocytometer and then stained in FACS buffer (0.5% BSA, 0.1 mM EDTA pH 8 and 1X PBS) with the following antibodies: anti-CD45 (clone 30-F11), anti-CD11c (clone N418), anti-CD19 (clone 1D3), anti-NK1.1 (clone PK 136), anti-CD25 (clone PC 61.5), anti-CD44 (clone IM7), anti-CD45R (cloneRA3-6B2), anti-CD64 (X54-5/7.1), anti-MerTK (clone D55MMER), anti-Ly6G (clone1A8), anti-SiglecF (clone E50-2440), anti-major histocompatibility complex (MHC) class II (clone M5/114.15.2), anti-CD11b (clone M1/70), anti-Ly6C (clone AL-21), anti-CD8 (clone 53-6.7), anti-CD3e (clone 17A2), anti-CD103 (clone 53-7.3), anti-CD4 (clone GK1.5), anti-TREM2 (clone 78.18). Antibodies were obtained from Cytek, Biolegend, BD, and eBioscience as previously described (68). Stained cells were run on an LSRFortessa cell analyzer (BD), and the data were analyzed using FlowJo software v10.10 (BD Biosciences). Gating strategies were used as previously described (68).

### Histopathology

After mice were euthanized, the lungs were perfused with 4% paraformaldehyde (PFA) in 1X PBS *in situ* with a catheter inserted through an incision made on the trachea.

Next, the lungs were removed, fixed in 4% PFA overnight, and then stored in 70% ethanol until processing. Lungs were embedded in paraffin and cut into 5 μm sections by the Cedars-Sinai Biobank and Research Pathology Resource (BRPR). The resulting slides were stained with hematoxylin and eosin (H&E) (Millipore Sigma). For further image analysis, slides were scanned using an Aperio AT2 slide scanner (Leica) in the BRPR. The scanned images were examined using Aperio Image Scope software (Leica); intracellular and extracellular *Cryptococcus* cells and macrophages from all fields of 2 lung slices per mouse were manually counted to determine the ratio of intracellular to total fungal cells, phagocytic index (total number of engulfed fungal cells per 100 macrophages), and percentage of uptake by macrophages (ratio of macrophages containing ≥1 fungal cell to total macrophages x 100).

### Cytokine measurements

Whole lungs from infected mice were homogenized in 2 mL PBS containing 1X Halt^TM^ Protease Inhibitor Cocktail using gentleMACS^TM^ C tubes with the “RNA 1” program on a gentleMACS^TM^ Octo Dissociator (Miltenyi Biotec). The samples were centrifuged at 1500 rpm to isolate the supernatant. Cytokines were quantified using the Bio-Plex Pro Mouse Cytokine 23-Plex Assay (Bio-Rad) following the manufacturer’s protocol on a Bio-Plex 200 System (Bio-Rad). The measured cytokines include IL-1α, IL-1β, IL-2, IL-3, IL-4, IL-5, IL-6, IL-9, IL-10, IL-12p40, IL-12p70, IL-13, IL-17, IFN-γ, TNF-α, MCP-1, Eotaxin, G-CSF, GM-CSF, KC, MIP-1α, MIP-1β, and RANTES.

### TREM2 binding study

TREM2-Fc binding to fungal cells was evaluated as previously described (14, 53). Briefly, recombinant TREM2-Fc protein that fuses the murine or human TREM2 extracellular domain with the Fc domain of human IgG (R&D 1729-T2 and 1828-T2) and purified human IgG (R&D 1-001-A), as a control, were incubated with PE-conjugated F(ab’)_2_ Fragment directed against the Fc domain of human IgG (Jackson ImmunoResearch 09-116-170) for 30 min at 4°C. The resulting antibody complexes were incubated with fungal cells for 1 h at 4°C. Samples were washed and resuspended in FACS buffer. Fungal cells bound to TREM2-Fc or human IgG were detected by flow cytometry using an LSRFortessa cell analyzer (BD) and FlowJo software (Waters Biosciences).

### Statistical analysis

All data are expressed as mean ± SEM. A *t*-test was used for statistical analysis of two group comparisons and one-way ANOVA was employed for 3 or more groups unless otherwise stated. Survival data were analyzed by Mantel-Cox test. All statistical analyses were carried out with GraphPad Prism software, v10.20. A *P* value < 0.05 was considered significant and indicated with an asterisk.

## Acknowledgments

We thank Jessica Hamerman, Fayyaz Sutterwala, Tamara Doering, Jennifer Lodge, Rajendra Upadhya, Camaron Hole, Dolores Di Vizio, and our colleagues in the Women’s Guild Lung Institute and Research Division of Immunology for reagents, equipment, and helpful discussions. We acknowledge assistance from the Cedars-Sinai Flow Cytometry Shared Resource and Biobank and Research Pathology Resource.

This work was funded in part by NIH/NIAID K08AI130366 to L.J.H and NIH/NIAID R01AI162765 to L.J.H. A.R. is supported in part by NIH/NHLBI T32HL170963. The funders had no role in the study design, data collection and interpretation, or the decision to submit the work for publication.

## Notes

### Competing Interest Statement

The authors have declared no competing interest.

### Summary of Updates

The supplemental files have been added to the manuscript.

