## Supplementary Material for "TREM2 drives monocyte-derived macrophage responses to *Cryptococcus neoformans*"

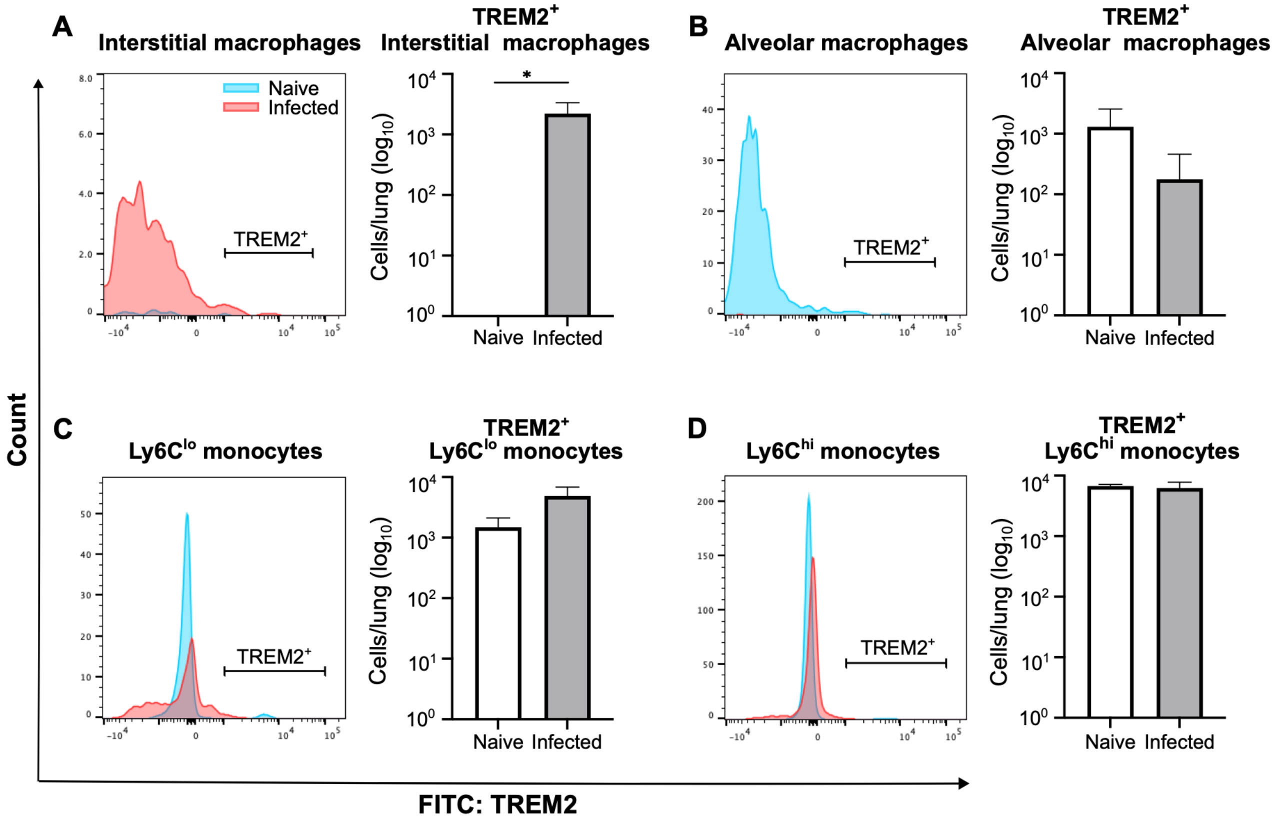


**Fig. S1: TREM2 expression by lung monocytes and macrophages on Day 21 p.i.**
(A-D) TREM2 expression on monocytes and macrophages from the lungs of naive and infected WT mice on Day 21 p.i. were analyzed via flow cytometry. Representative histograms are shown on the left of each panel. Blue = naive, Pink = infected. Quantification of the number of $TREM2^+$ cells in each subpopulation is shown in the bar graphs on the right of each panel. White = naive, Gray = infected. Data are from one independent experiment (*n* = 5 total mice per group). *, $P < 0.05$ by *t*-test.

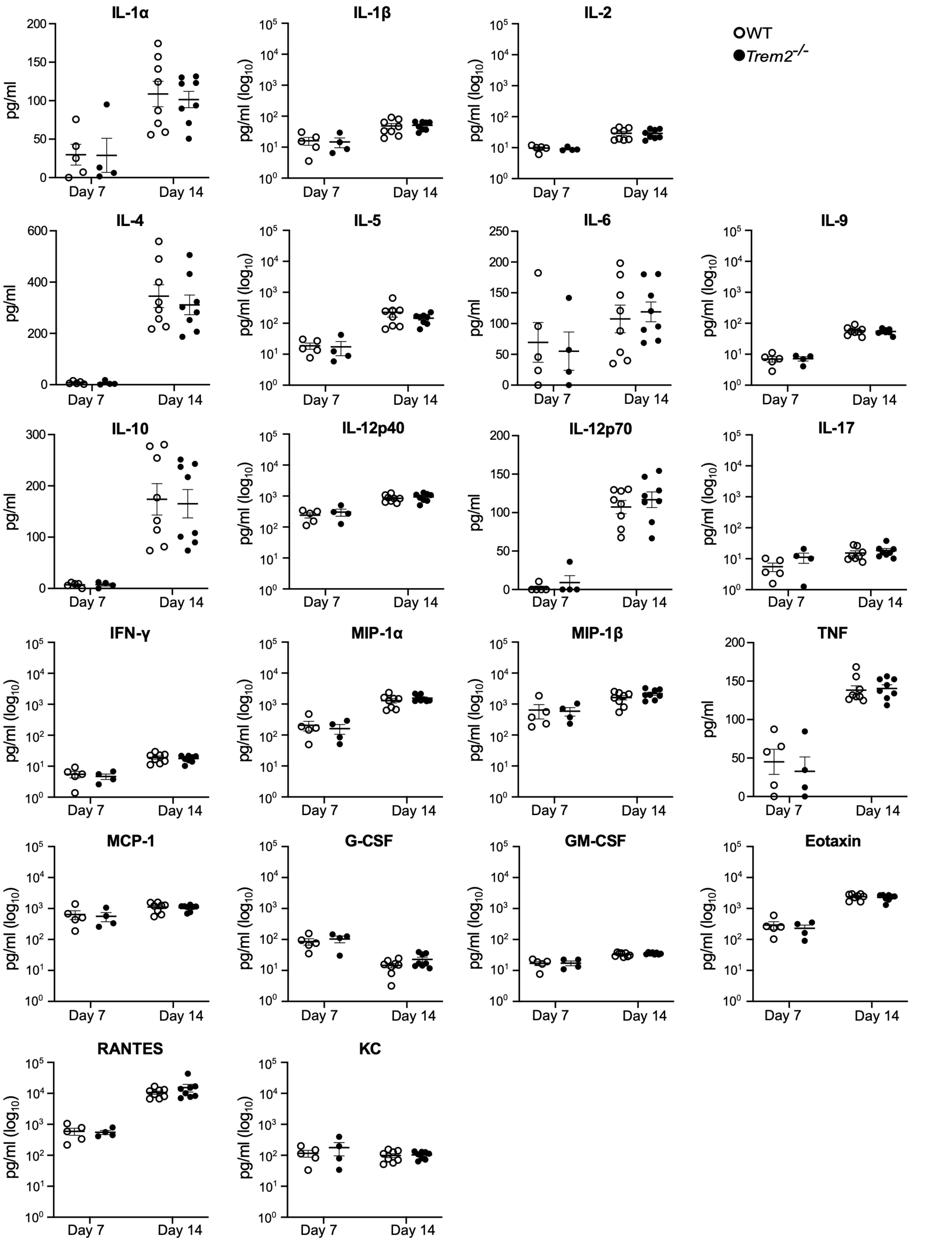


**Fig. S2: Lung cytokine responses are not altered by TREM2 after *C. neoformans* infection**. Cytokine levels in the supernatant of whole lung homogenates from WT and *Trem2*$^{-/-}$ mice on Days 7 and 14 p.i. with *C. neoformans.* Data are pooled from two independent experiments ($n$ = 7-9 total mice per group) and analyzed by $t$-test.

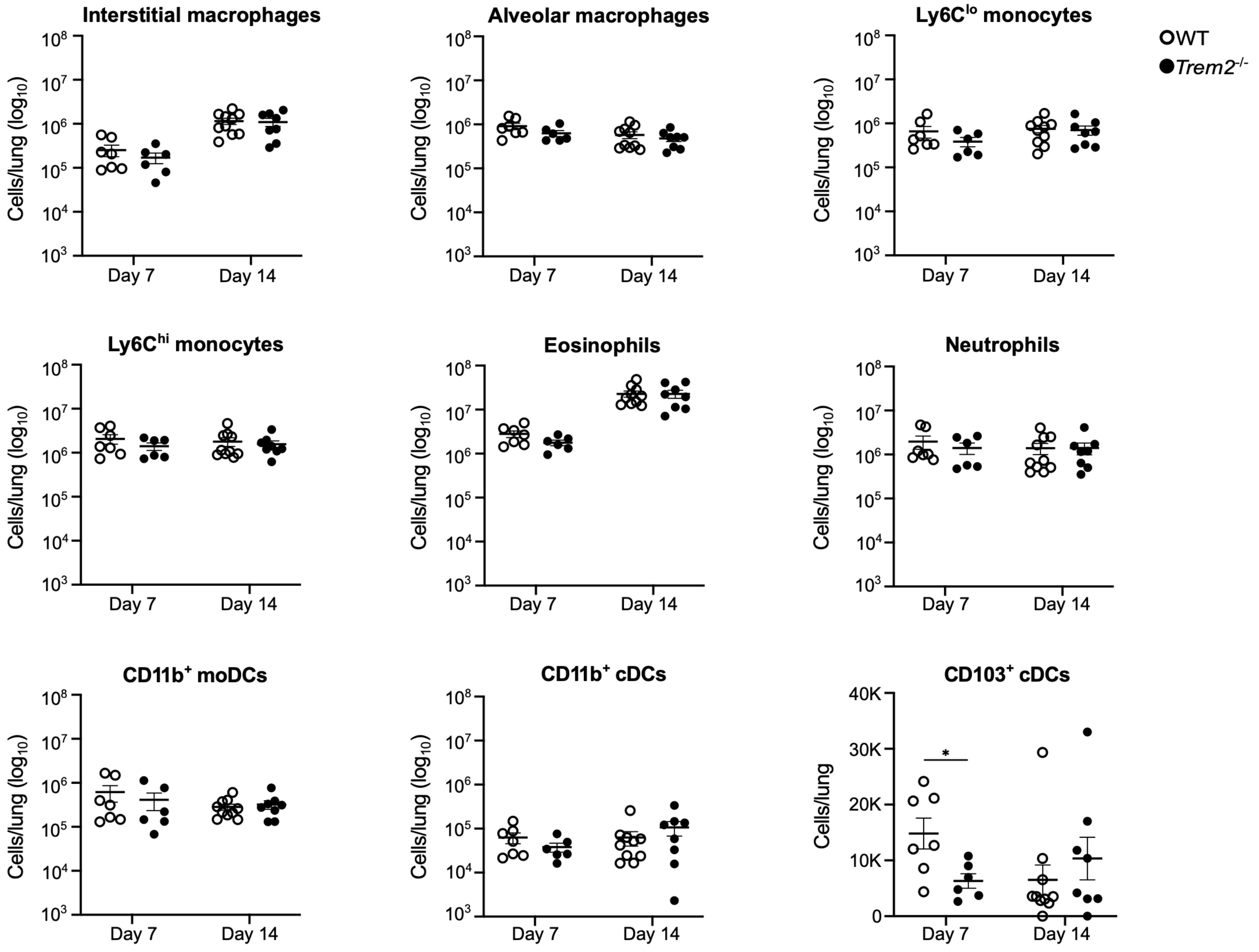


**Fig. S3: TREM2 does not regulate the recruitment of innate immune cells populations to the lungs during infection with *C. neoformans***. Immune cells from the lungs of WT and *Trem2*$^{-/-}$ mice on Days 7 and 14 p.i. were analyzed by flow cytometry. Monocyte-derived dendritic cells (moDCs), conventional dendritic cells (cDCs). Data are pooled from two independent experiments ($n$ = 6-7 total mice per group). *, $P < 0.05$ by $t$-test.

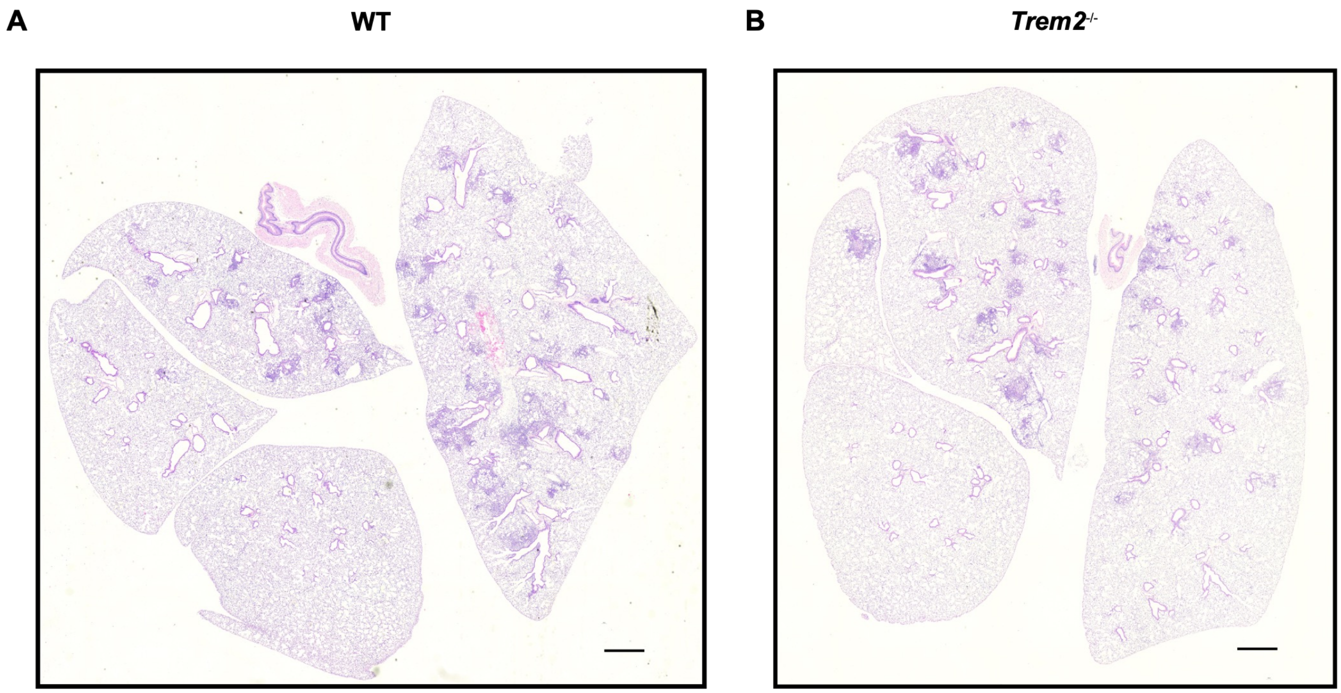


**Fig. S4: TREM2 does not affect overall lung pathology during *C. neoformans* infection.** Representative images of whole lung sections from (A) WT and (B) *Trem2*$^{-/-}$ mice on Day 7 p.i. stained with H&E (*n* = 3 total mice per group). Scale bars = 1000 μM at 4X magnification.

| Fungal strains | Parental strain | Drug selection | Reference |
|---|---|---|---|
| *ags1*Δ | H99R | URA5 | (32) |
| *cda1Δcda2*Δ | KN99 | HYG/NAT | (35) |
| *cda1Δcda2Δcda3*Δ | KN99 | HYG/NAT/Phleo | (35) |
| *chs3*Δ | KN99 | NAT | (69) |
| *kre5*Δ | H99 | HYG | (37) |
| *skn1*Δ*kre6*Δ | H99 | HYG/NAT | (37) |

**Table S1: *C. neoformans* mutant strains used in this study.**
